# A diffusion model conditioned on compound bioactivity profiles for predicting high-content images

**DOI:** 10.1101/2024.10.10.616543

**Authors:** Steven Cook, Jason Chyba, Laura Gresoro, Doug Quackenbush, Minhua Qiu, Peter Kutchukian, Eric J. Martin, Peter Skewes-Cox, William J. Godinez

## Abstract

High-content imaging (HCI) provides a rich snapshot of compound-induced phenotypic outcomes that augment our understanding of compound mechanisms in cellular systems. Generative imaging models for HCI provide a route towards anticipating the phenotypic outcomes of chemical perturbations in silico at unprecedented scale and speed. Here, we developed Profile-Diffusion (pDIFF), a generative method leveraging a latent diffusion model conditioned on in silico bioactivity profiles to predict high-content images displaying the cellular outcomes induced by compound treatment. We trained and evaluated a pDIFF model using high-content images from a Cell Painting assay profiling 3750 molecules with corresponding in silico bioactivity profiles. Using a realistic held-out set, we demonstrate that pDIFF provides improved predictions of phenotypic responses of compounds with low chemical similarity to compounds in the training set compared to generative models trained on chemical fingerprints only. In a virtual hit expansion scenario, pDIFF yielded significantly improved expansion outcomes, thus showcasing the potential of the methodology to speed up and improve the search for novel phenotypically active molecules.

## 1 Introduction

Compound-induced phenotypes observed through high-content imaging (HCI) assays provide rich clues as to the activity and mechanisms of compounds in cellular systems (Bray et al, 2016). While HCI assays can screen large compound collections, experimental assays are still restricted to a comparatively small sample of the synthetically accessible chemical space (Dobson, 2004; Drew et al, 2012; Grygorenko et al, 2020). Generative models that predict images containing the specific phenotype induced by any compound can enable virtual screening and profiling of compound collections at unprecedented scale, speed, and cost (Yang et al, 2021).

Early work in generative models for cell microscopy images focused on mapping a symbolic representation of cellular biology (e.g., a state vector) onto an image using explicit appearance models for image generation (Murphy, 2005; Zhao and Murphy, 2007; D. V. Sorokin et al, 2018). More recent approaches leverage advances in deep neural networks (LeCun et al, 2015) to learn image generation functions directly from training images (Osokin et al, 2017; Johnson et al, 2017; Goldsborough et al, 2017). Conditioning the generation functions with compound representations, such as compound fingerprints, allows these methods to generate high-content images showing the phenotypic outcomes induced by compound treatment (Yang et al, 2021; Palma et al, 2023). Architecturally, these methods are built upon flow-based generative models (Yang et al, 2021) and generative adversarial networks (GANs) (Palma et al, 2023).

Recent research points to the superior efficacy of diffusion−based generative architectures for conditional image generation (Dhariwal and Nichol, 2021), and more broadly, for drug discovery tasks (Abramson et al, 2024; Corso et al, 2023). Chemically, these methods are conditioned on compound representations derived directly from molecular structures, limiting model generalizability to structurally similar chemical matter (Yang et al, 2021; Palma et al, 2023). Alternative representations based on imputed compound bioactivity (Martin et al, 2019) can potentially improve the ability of such generative methods to extrapolate to novel chemical matter.

In this paper, we introduce Profile Diffusion (pDIFF), an approach combining a stable diffusion-based generative model with in silico bioactivity profiles to predict high-content images displaying the cellular outcomes induced by compound treatment. We build and validate a pDIFF model with a collection of bioactivity profiles and Cell Painting images corresponding to a diverse chemogenomic library of 3750 compounds. Using a “realistically novel” split for validation (Martin et al, 2019), we demonstrate that pDIFF improved extrapolation to novel chemical matter compared to a baseline diffusion model conditioned on chemical fingerprints. We also test the pDIFF model in a virtual hit expansion scenario and show that pDIFF yields significantly improved hit expansion outcomes.

## 2 Results

### 2.1 Profile Diffusion: Stable diffusion model conditioned on compound profiles

We developed the Profile Diffusion (pDIFF) methodology based off the stable diffusion architecture (Rombach et al, 2022). At training time, pDIFF takes as input an in silico bioactivity profile as well as a high-content image displaying the cellular outcomes induced by that compound. The input image is compressed into a lower-resolution latent representation through a pre-trained variational autoencoder (VAE) (Kingma and Welling, 2022). A latent diffusion network conditioned on the compound’s bioactivity profile is trained in the VAE’s latent space to predict the noise added to the latent image at each step of a diffusion process (see **Methods**). Thus, in comparison with the original implementation of the Stable Diffusion model, where natural language embeddings are used to condition the denoising network, pDIFF instead induces the model to learn the “language of compound bioactivity”.

To generate an image for a specific compound with a trained pDIFF model, we take the in silico bioactivity profile of that compound, sample a random noise tensor in the VAE’s latent space, and iteratively denoise that tensor with the model conditioned on the compound’s bioactivity profile together with a denoising diffusion probabilistic models (DDPM) schedule (Ho et al, 2020) (see **Methods**). Varying the number of denoising iterations trades off image quality against computation time, as the model must be invoked more often for more steps. Finally, the denoised tensor is passed through the VAE decoder to synthesize a high-content image. Different random noise tensors result in different images, thus allowing generation of a collection of images for each input compound.

### 2.2 Profile Diffusion generalizes to novel chemical matter

To validate pDIFF in a realistic drug discovery context, we used 3750 compounds of the Novartis Mechanism-of-Action (MoA) Box chemogenomic library (MoA Box) (Canham et al, 2020). These compounds were profiled in a Cell Painting assay (Bray et al, 2016), where U-2 OS cells were compound-treated in triplicate at 12.5 *µ*M (see **Methods**). Twenty-four high-content images were acquired per compound. We utilize the cytoplasm, mitochondria, and nuclei channels of the Cell Painting protocol for model building and evaluation.

As conditioning profile for pDIFF, we used the in silico bioactivity profiles predicted by Profile-QSAR (pQSAR) (Martin et al, 2019), a massively multitask bioactivity machine learning model trained on over two million compounds across 14222 Novartis dose-response assays (see **Methods**). For each of the 3750 compounds in our dataset, pQSAR predicts activity in terms of pAC_50_ values (which is the negative logarithm of the half-maximal activity concentration) for each of the 14222 assays, thus resulting in a 14222-length profile per compound. For numerical tractability, we reduced predictions from the 14222 assays to 2018 assays through a target-focused assay selection scheme (see **Methods**).

For benchmarking purposes we also trained a stable diffusion model conditioned on chemical fingerprints. Specifically, we used extended-connectivity fingerprints (ECFPs) (Rogers and Hahn, 2010), each folded into a 2048 count vector. For this baseline model, we used the same training and inference regimes as used for pDIFF. At inference time, we set both pDIFF and the baseline model to generate 12 images per compound (see **Methods**).

To validate the performance of pDIFF and the baseline model, we used a “realistically novel” cluster-based split approach (Martin et al, 2019). This approach aims to replicate a realistic screening scenario, where project teams are interested in testing compounds that differ significantly from those previously tested (see **Methods**). We chose 10% of the total dataset size for the held-out set, giving us 3375 training and 375 held-out compounds. This split results in a highly dissimilar median Tanimoto coefficient of 0.11 between train and test compounds (**Supplementary Figure 1**).

Example real and generated images for three held-out compounds are shown in **Figure 2**. These active compounds induce outcomes visually different from those observed in the neutral controls (see **Supplementary Figure 2**). We show images for Halofuginone, which is known to inhibit the viability of Osteosarcoma cell lines (Lamora et al, 2015). The baseline model conditioned on chemical fingerprints is unable to correctly predict the cell death outcome induced by this compound, which has a low chemical similarity to the training set (Tanimoto coefficient of 0.23 to the nearest neighbor compound in the training set). In contrast, the pDIFF model, which is conditioned on pQSAR bioactivity profiles, visually recapitulates this outcome. We also show images for CHEMBL2326002, a protein kinase C-theta inhibitor which induces a slightly elongated phenotype with a punctuated pattern in the mitochondria channel. The baseline model does not predict the elongated punctuated pattern, whereas pDIFF generates images with cells closely resembling the phenotype induced by this compound. For Brusatol, which plays a role in DNA damage repair (Li et al, 2023) by acting as an inhibitor of the NRF2 pathway (Ren et al, 2011), cells exhibit a toxic phenotype that is anticipated only by pDIFF.

**Fig. 1.**
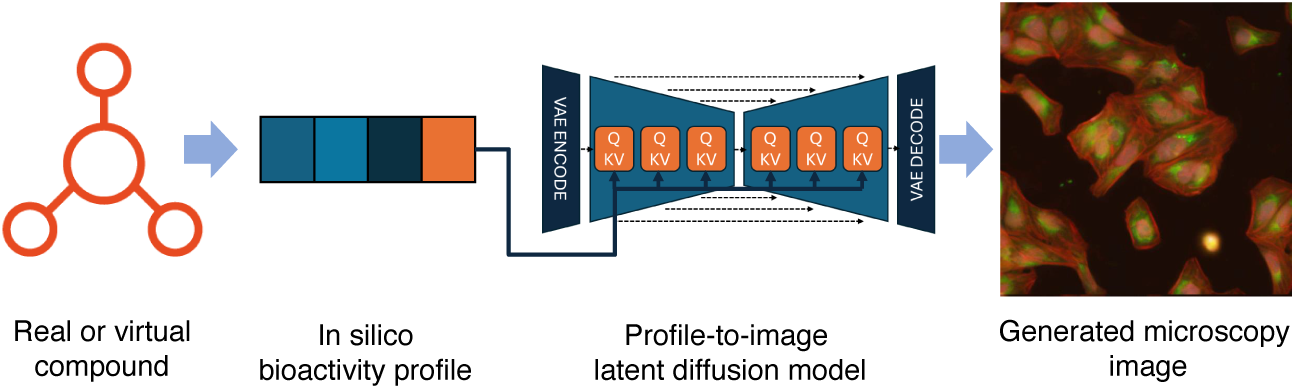
Profile-Diffusion (pDIFF) workflow. A virtual or real compound is represented computationally through an in silico bioactivity profile computed by pQSAR. The representation is used to condition the generative process of the stable diffusion model underlying pDIFF to generate a high-content image showing the phenotypic outcome induced by compound treatment.

**Fig. 2.**
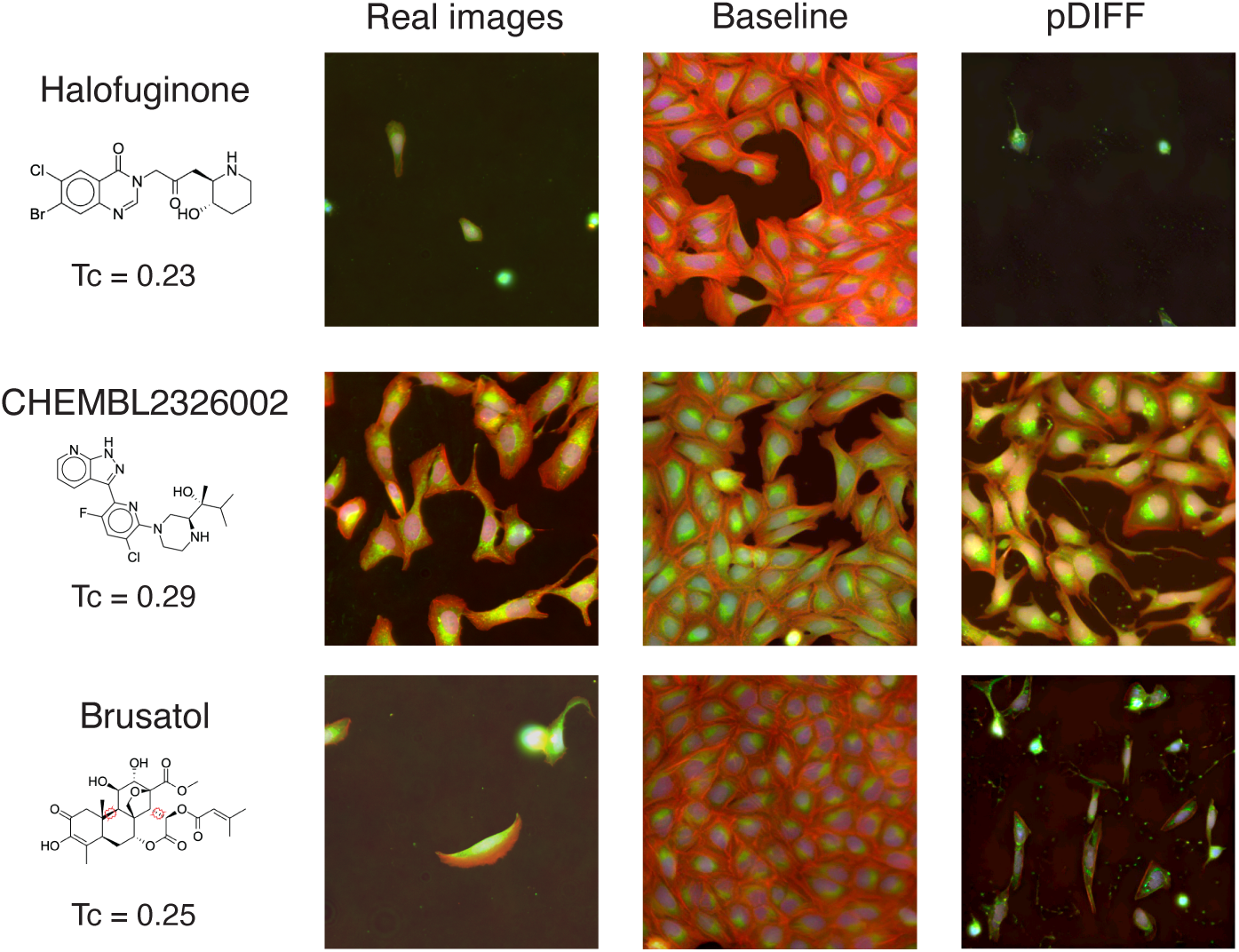
Molecules in the realistic held-out set along with their corresponding real images (Nuclei, blue; mitochondria, green; red, cytoplasm) as well as images generated with diffusion models. We show images generated by a baseline diffusion model conditioned on chemical fingerprints (Baseline) as well as by pDIFF. For each molecule, we list the Tanimoto coefficient (Tc) to the nearest neighbor molecule in the training set.

**Table 1.** Results for held-out validation compounds. Values given are Spearman correlation coefficients between compound-aggregated average values for each image feature. The first row shows the correlation values for each half of the real images compared against each other. pDIFF model performs better than the baseline diffusion model conditioned on chemical fingerprints.

|  | Coverage | Cell Count | Cell Size |
| --- | --- | --- | --- |
| Real images upper bound | .67 | .56 | .48 |
| Baseline diffusion model | .13 | .10 | .04 |
| pDIFF | .48 | .28 | .12 |

To quantitatively assess the ability of pDIFF to recapitulate phenotypic outcomes, we follow a previous scheme (Yang et al, 2021) by collecting image- and cell-level features (viz. coverage, cell count, and cell size) for real and generated image sets. Features are averaged over all images for each compound, then Spearman correlation coefficients between these compound-aggregated features of real and generated image sets are calculated (see **Methods**).

**Table** 1 shows the real vs. generated correlation coefficients across the 375 held-out (test) compounds for each of the six features. As upper bound, we compute the correlation among real images of the held-out compounds by splitting the 24 real images into two halves, calculating features for each half, and computing the correlation coefficients between the features of both halves. Correlation values for the image features between the two sets of real images range from 0.48 to 0.67. For this challenging held-out set of compounds with low chemical similarity to the training set, the baseline diffusion model conditioned on chemical fingerprints yielded correlation values ranging from 0.04 to 0.13. The pDIFF model yielded correlation coefficients between 0.12 to 0.48, thus exhibiting substantially improved performance on novel chemical matter compared to the baseline model.

### 2.3 Profile Diffusion enables improved retrieval of phenotypically similar molecules for hit expansion

Having ascertained the ability of pDIFF to generalize to novel chemical matter, we proceeded to test pDIFF in a virtual hit expansion scenario, where the goal is to find compounds inducing phenotypic outcomes similar to those induced by known active compounds (i.e. phenotypic screening hits). To set the ground truth for this scenario, we take the real images of 101 active query compounds from the training set and compare these to the real images of the 375 compounds in the held-out (test) set. To compare the similarity of two compounds via images, we use the Sinkhorn divergence (Feydy et al, 2019) on the sets of Cellpose (Stringer et al, 2021) feature vectors derived from the compounds’ corresponding images. Using these Sinkhorn divergences, we retrieve the 50 nearest-neighbor compounds in the test set for each query compound in the training set. We repeat the nearest neighbor retrieval but with pDIFF-generated images instead of real images for the test compounds, thus comparing real images for query compounds with pDIFF synthetic images for test compounds. As performance measure for this task, we calculate the percentage overlap between the ground truth nearest neighbors and those retrieved using pDIFF images.

We report the distribution of percentage overlap between the real and pDIFF test-set nearest-neighbors for the 101 query compounds in **Figure** 3. As baseline, we show the percentage overlap values for 101 random selections of 50 test compounds, which yields a median percentage overlap of 14%. As additional baselines, we show the percentage overlap values for nearest neighbors retrieved with chemical fingerprints as well as bioactivity profiles (ECFP and pQSAR profiles, respectively) using a Tanimoto or cosine distance. These approaches lead to median percentage overlaps of 16% and 38%, respectively. A baseline diffusion model conditioned on ECFPs leads to a median percentage overlap of 16%. Finally, retrieval with images generated by pDIFF leads to a median percentage overlap of 50%. A two-sample Kolmogorov–Smirnov test (Conover, 1971) was used to determine the statistical significance of differences among the percentage overlap values of the different approaches. The resulting p-values were corrected for false discovery rate with the Benjamini–Hochberg method (see **Supplementary Table 1**). The differences between the distribution of percentage overlap values of the pDIFF approach and those of all other approaches were significant. This outcome suggests that pDIFF enables improved retrieval of phenotypically similar molecules, thus enhancing virtual expansion outcomes.

**Fig. 3.**
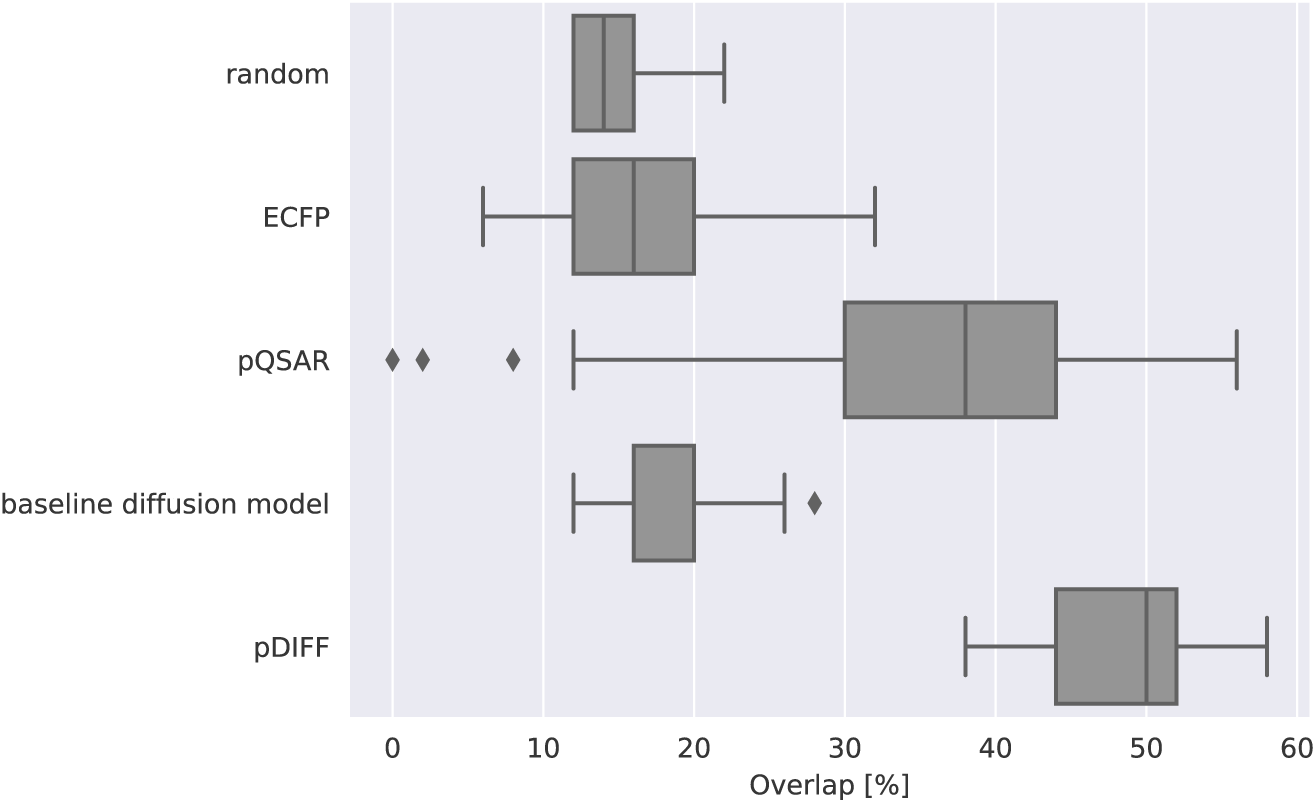
Distribution of percentage overlap values between 50 ground truth nearest neighbors and those retrieved with virtual expansion approaches for 101 query compounds. We show the performance for an approach selecting molecules randomly (‘random’) as well as for similarities computed using chemical or bioactivity profiles (ECFP and pQSAR, respectively). The results for approaches computing similarities using image profiles derived from images generated through either a baseline diffusion model or pDIFF are also shown.

## 3 Discussion

High-content screening assays provide rich mechanistic and holistic information about cellular responses to chemical perturbations. Overcoming fundamental limitations in screening capacity and compound synthesis via in silico generated high-content images holds potential to vastly expand the compound search space. To this end, we developed a purely virtual generative imaging approach, pDIFF, necessitating only in silico bioactivity profiles as input for image generation. Using a realistic validation scheme, we show that pDIFF provides remarkably improved image predictions for novel chemical matter.

Generating HCI images instead of predicting image measurements directly from compound representation benefits from state-of-the-art diffusion models that both excel at generating high-quality, high-dimensional images as well as exhibiting more stable convergence (Huang et al, 2024). Relevant image-derived measurements reflective of phenotype can subsequently be extracted from the generated images with existing image analysis pipelines. In our study, for example, we use the same analysis pipeline to segment and extract image features for both real and pDIFF-generated images. While the image features could be predicted directly from in silico compound profiles, the ability to generate images is key to gaining the confidence of experimentalists as the generated images readily provide visual insight informing the image-derived measurements. pDIFF allows for visual review of HCI-based virtual screens just as chemists visually review virtual screens from chemical similarity or docking.

In our study, we used the bioactivity profiles calculated by the in-house Profile-QSAR (pQSAR) model (Martin et al, 2019) trained on over 2M compounds and 14k Novartis internal dose-response assays. The pQSAR methodology is publicly available, and useful models have been built on public bioactivity databases, such as ChEMBL (Zdrazil et al, 2023). pDIFF could also be trained on other experimental profiles such as gene expression signatures (Ye et al, 2018; Li et al, 2012; Subramanian et al, 2017). Furthermore, augmenting the bioactivity profiles with cell type and concentration information with corresponding image data could expand the predictive scope of a single pDIFF model.

Given its ability to predict phenotypic outcomes for novel chemical matter, we envision pDIFF having impact in a number of drug discovery scenarios. As shown by our findings, hit expansion campaigns searching for molecules inducing phenotypic outcomes similar to those induced by hit molecules can benefit from the improved image-based similarity calculations enabled by pDIFF. Profiling activities (Chandrasekaran et al, 2021) aiming to reveal the mechanism-of-action of molecules can likewise leverage pDIFF generated images for similarity calculations relative to images of a reference set of annotated molecules. Image-based readouts have been shown to be predictive of cardiotoxocity (Garcia de Lomana et al, 2023; Seal et al, 2024) and we anticipate pDIFF-generated images to help flag adverse cardiac effects of compounds. Historical imaging assays could be revisited virtually through pDIFF, so past image screening investments can be leveraged to enable future follow up work.

Training pDIFF is computationally demanding due to the need to re-learn the space of high-content images and their corresponding profiles instead of the CLIP text embeddings Stable Diffusion was trained with. Our training set of 12 fields of view (FOVs) per each of 3,375 compounds, took 360 GPU-hours to train for 30,000 steps.

A trained pDIFF model can predict one 512×512 pixel output patch per GPU-second using a conservative, high-quality inferencing schedule (DDPM (Ho et al, 2020), 100 network inference steps). Alternative schedulers such as the Diffusion Probabilistic Models (DPM) solver (Lu et al, 2022) purport to give a similar quality of results using only tens of steps, making this an attractive area for further optimization. Diffusion model inference is more expensive than other generative models like GANs due to the need to repeatedly invoke the network. Even with our conservative settings, the entire MoA box imageset of 90,000 images (3,750 compounds × 24 FOVs) could be re-generated via inference in 25 GPU-hours, without need for expensive and time-consuming compound synthesis, plating, image acquisition, etc.

In conclusion, our work shows the potential of in silico image prediction to anticipate induced phenotypic outcomes of compounds, including those generated through other machine learning methods (Sanchez-Lengeling and Aspuru-Guzik, 2018; Godinez et al, 2022; Shen et al, 2024), much earlier in the drug discovery pipeline. The application of pDIFF allows us to explore broader and more diverse compound collections at unprecedented speed, scale, and efficiency.

## Acknowledgements

We thank Frederick Lo for help with data logistics and pre-processing. We thank Mark A. Bray for fruitful discussions.

## Data availability

The data used in this study is proprietary to Novartis. The data is not publicly available due to intellectual property restrictions. An example dataset is available in the pDIFF code respository.

## Code availability

The code and an example dataset for pDIFF is available in **Supplementary Code** and at https://github.com/Novartis/pDIFF ^1^.

## Author contribution

S.C and W.J.G. designed and led the study. S.C. developed, implemented, and evaluated pDIFF. J.C, L.G., and D.Q. developed and ran the imaging assay. E.J.M. developed the algorithm to compute the in silico bioactivity profiles and provided feedback. M.Q., P.K., and P.S.-C. provided feedback. S.C., M.Q., and W.J.G. analyzed and interpreted the results. S.C. and W.J.G. wrote the article. All authors reviewed the manuscript.

## 7 Methods

### 7.1 Image Acquisition and Preparation

U2-OS cells were treated with 3750 compounds from the Novartis MoA Box (Canham et al, 2020) at a fixed concentration of 12.5µM and incubated for 24 hours. The Cell Painting staining protocol (Bray et al, 2016) was applied. Image acquisition was performed using a 4 HP laser, 4 camera Phenix imaging system and a 20x NA1.0 water immersion objective. Acquired images were background corrected using the BaSIC algorithm (Peng et al, 2017), rescaled by a factor of ½ from 2160x2160 pixels to 1080x1080, then random crops of 512x512 are extracted from each image for training. RGB images were assembled from the F-actin Cytoskeleton, Mitochondria, and Nucleus channels (Phalloidin/Alexa, MitoTracker Deep Red, and Hoescht 33342 stainings, respectively).

### 7.2 Profile calculations

To compute the in silico bioactivity profiles for each compound, we used 14222 models from the massively-multitask pQSAR algorithm for predicting pAC_50_ values for 14222 Novartis-internal biochemical and cellular dose-response assays. We narrowed the resulting 14222-D bioactivity profile by first selecting only the assays where pQSAR models provided useful predictions on the challenging ”realistically novel” held-out set per assay, as measured by the squared Pearson correlation coefficient *r*^2^ *>* 0.3 between predicted and experimental pAC_50_ values. We then selected the target-based biochemical and cellular assays in the narrowed assay collection, grouped them by target protein, and selected the best-performing pQSAR model based on the models’ *r*^2^ values per target. For the remaining purely-phenotypic assays, we grouped them by drug discovery project, and likewise selected the best-performing pQSAR model per project. For the remaining set of assays with neither target nor project annotations, we selected only very high-quality pQSAR models (*r*^2^ *>* 0.9). This selection process resulted 2018 assays, amounting to a 2018-D in silico bioactivity profile per compound. The profile was zero-padded to reach the 2048-D input required into the pDIFF model.

### 7.3 Model Architecture

The Stable Diffusion 2.1 model (Rombach et al, 2022) was used as the backbone for pDIFF. The lengths of the cross-attention weights are changed to 2048, and all existing weights are re-initialized. As no natural language inputs are needed, the CLIP module is removed. The pre-trained VAE is retained and its weights are frozen. Following (Rombach et al, 2022), we train the model to minimize the mean squared error between actual noise *ɛ* and predicted noise *ɛ_θ_*(*z_t_, t, y*) (Eq. 1), where *t* is a current timestep of the noise schedule, *θ* are the network parameters, *z_t_* is the latent image representation at step *t* of the noise schedule, E is the VAE used to encode the image *x*, and *c* is the conditioning profile.

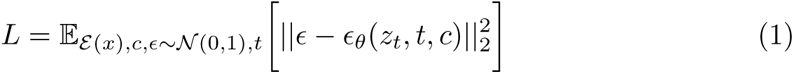

### 7.4 Model Training

pDIFF was trained for 30k steps on 4x Nvidia A100 GPUs using huggingface Accelerate, no mixed precision, in distributed data parallel mode. pDIFF is trained on 12 images per training set compound, as the remaining 12 are reserved for calculating an upper bound for expected image similarity. Training time was approximately 90 wall-clock hours (360 GPU hours) for the 40k images in the training set (12 images per compound x 3375 compounds). The Min-SNR weighting strategy (Hang et al, 2023) was applied to accelerate convergence, and offset noise (Guttenberg, 2023) was applied to help the model learn to generate empty and nearly-empty FOV images with very low average intensity. Training hyperparameters are presented in **Supplementary Table 2**.

### 7.5 Inference

The DDPM scheduler (Ho et al, 2020) with 100 steps was used for inference with the model. 12 images were generated per compound profile, at a time cost of approximately 1 second per 512x512 output image per GPU. We use the guidance algorithm (Ho and Salimans, 2022) to drive the generation process. In Eq. 2 *w* is the guidance scale, *c* is the conditioning profile, *z_t_* is the output result of the previous denoising step, ø is a null conditioning profile, *ɛ_t_*(*z_t_,* ø) is the model’s unconditional noise prediction, *ɛ_t_*(*z_t_, c*) is the conditional noise prediction, and *ɛ*~*_t_* is the resulting guided noise (Ho and Salimans, 2022).

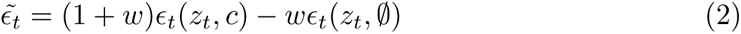

Inference-time classifier-free guidance scale value *w* = 4 was chosen empirically.

### 7.6 Model Validation

We used the “realistically novel” split (Martin et al, 2019) to evaluate pDIFF. In this approach, the compounds are clustered by similarity, and the clusters ranked in order of size. We allocated compounds from the larger clusters to the training set until 3375 (90%) of the compounds were included. The remaining 375 (10%) of compounds coming from the singletons and smaller clusters were allocated to the realistic held-out set.

To quantify model performance, we first segmented real and generated images using Cellpose with the default cyto2 model in grayscale mode (Stringer et al, 2021). Following a previous approach (Yang et al, 2021), hand-engineered features were calculated for each image. Specifically, we computed the total image area covered by segmented cells (coverage), the number of segmented cells in the image (cell count), and the size of the segmented cells. We averaged the values of these features over all images corresponding to the same compound and calculated Spearman correlations coefficients between features derived from real and generated images across the 375 held-out, or test, compounds.

### 7.7 Reproducing the Training Set

We check that models learned to reproduce phenotypic outcomes by evaluating the performance on the training set. **Supplementary Table 3** shows the Spearman correlation coefficients comparing pDIFF generated images for the 3375 molecules in the training set to the real images. As upper bound, we report the correlation values computed between the 12 real images used for model building and the remaining 12 held-out real images of the training molecules. Correlation values among real images across the calculated features range from 0.44 to 0.62. A baseline diffusion model conditioned on chemical fingerprints yields correlation values between 0.15 to 0.41. The pDIFF model shows very realistic performance with correlation values ranging from 0.40 to 0.59.

## Footnotes

1 GitHub repository to be published upon manuscript acceptance

