## Supplementary Information for "A diffusion model conditioned on compound bioactivity profiles for predicting high-content images"

[1]Novartis Biomedical Research, San Diego, 92121, CA, USA.
[2]Novartis Biomedical Research, Cambridge, 02139, MA, USA.
[3]Novartis Biomedical Research, Emeryville, 94608, CA, USA.

### Supplementary Figures

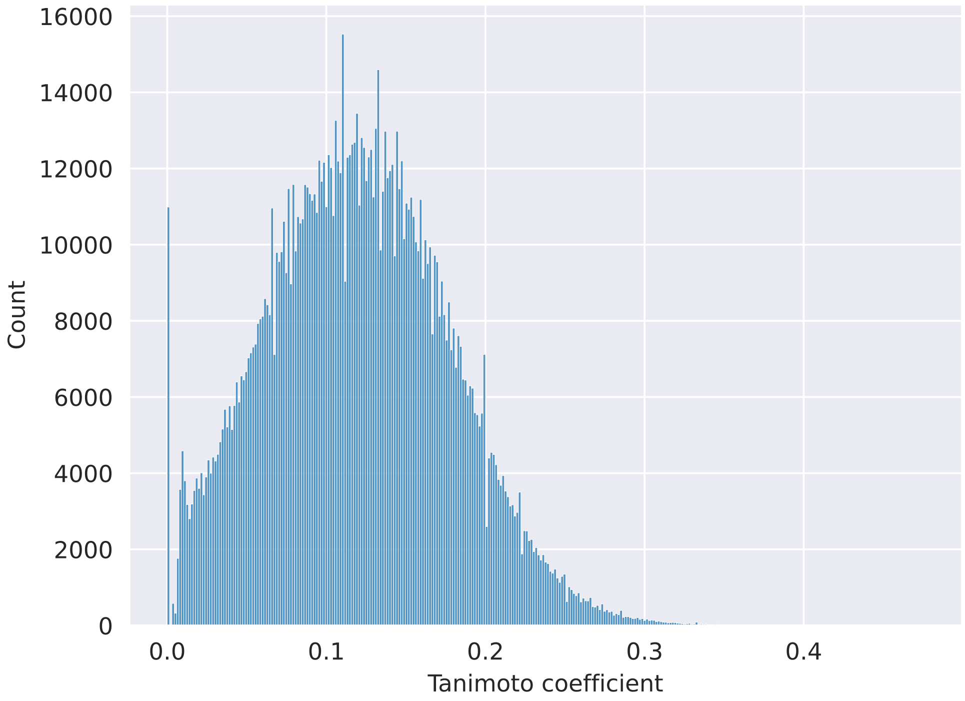


**Supplementary Figure 1** Histogram of Tanimoto coefficients between compounds in the training and held-out set for a total of 1265625 comparisons. The median Tanimoto coefficient is 0.11.

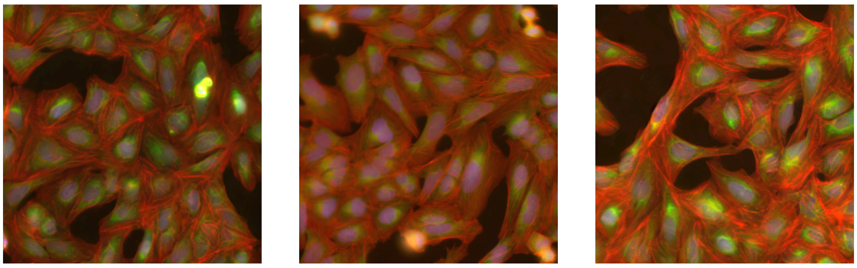

**Supplementary Figure 2** Examples images of the neutral control treatments (dimethyl sulfoxide, DMSO).

### Supplementary Tables

**Supplementary Table 1** Benjamini-Hochberg-adjusted p-values of two-sample Kolmogorov-Smirnov tests. Each test is carried out by comparing the distributions of percentage overlap values of two approaches.

| | random | ECFP | pQSAR | Baseline diffusion model | pDIFF |
|---|---|---|---|---|---|
| random | | 1.55e-03 | 3.45e-43 | 5.24e-07 | 1.85e-59 |
| ECFP | 1.55e-03 | | 5.45e-28 | 3.97e-03 | 1.85e-59 |
| pQSAR | 3.45e-43 | 5.45e-28 | | 9.19e-29 | 2.05e-13 |
| Baseline diffusion model | 5.24e-07 | 3.97e-03 | 9.19e-29 | | 1.85e-59 |
| pDIFF | 1.85e-59 | 1.85e-59 | 2.05e-13 | 1.85e-59 | |

**Supplementary Table 2** Training parameters for pDIFF.

| Parameter | Value |
|---|---|
| batch size | 6 |
| gradient accumulation steps | 10 |
| GPUs | 4x Nvidia A100 40GB |
| effective batch size | 240 |
| lr | 1e-4 |
| max grad norm | .1 |
| training steps | 30,000 |
| training schedule | constant, 1000 step warmup |
| noise offset | .2 |
| SNR gamma | 5.0 |
| optimizer | AdamW |
| adam beta1, beta2 | .9, .999 |
| adam weight decay | 1e-2 |
| adam epsilon | 1e-8 |

**Supplementary Table 3** Results for held-out images from training compounds. Values given are Spearman correlation coefficients between compound-aggregated average values for each image feature. The first row shows the correlation values between training images and held-out images.

| Configuration | Coverage | Cell Count | Cell Size |
|---|---|---|---|
| Real images upper bound | .62 | .53 | .44 |
| Baseline diffusion model | .41 | .36 | .15 |
| pDIFF | .59 | .46 | .40 |
